# Immune-mediated tumor control in the 5TGM1 transfer model of multiple myeloma

**DOI:** 10.1101/2023.01.18.524505

**Authors:** Zoltán Kellermayer, Sabrin Tahri, Madelon M. E. de Jong, Natalie Papazian, Cathelijne Fokkema, Remco Hoogenboezem, Mathijs A. Sanders, Louis Boon, Chelsea Den Hollander, Annemiek Broijl, Pieter Sonneveld, Tom Cupedo

**Author notes:** **Corresponding author** Tom Cupedo, Department of Hematology, Erasmus MC, Mailing address: Wytemaweg 80, 3015CN Rotterdam, The Netherlands.

## Abstract

Multiple myeloma is a disease of malignant plasma cells residing in the bone marrow, where interactions with local immune cells are thought to contribute to disease pathobiology. However, since a multiple myeloma diagnosis is virtually always preceded by an asymptomatic precursor phase, identifying early alterations in the bone marrow micro-environment following occupation by multiple myeloma cells remains challenging. Here we used the 5TGM1 transfer model of murine myeloma in combination with myeloma-permissive KaLwRij mice and myeloma-resistant C57Bl/6 mice and hypothesized that differential sensitivity to myeloma in these HLA-identical mouse strains has an immunological basis and might allow for dissection of early immune responses to myeloma cells.

Using flow cytometry and single-cell RNA sequencing we show that C57Bl/6 mice can restrain tumor growth for prolonged periods, associated with activation of cytotoxic immune responses that were absent from KaLwRij mice. Transcriptional analysis of immune cells and stromal cells identified a central role for IFN-signaling in tumor containment, and antibody-mediated neutralization of IFNγ increased both incidence and outgrowth of multiple myeloma in C57Bl/6 mice. Together these findings highlight the ability of a fully functional immune system to control multiple myeloma progression in an IFNγ−dependent manner and suggest that transfer of 5TGM1 cells into parental C57Bl/6 mice can serve as a faithful model to track anti-myeloma immune responses in immune competent and genetically modifiable mice.

## Introduction

Multiple myeloma (MM) is the second most common hematological cancer in adults and is characterized by accumulation of malignant plasma cells in the bone marrow (BM) (1,2). Recent advances in treatment options including the introduction of immunotherapies have significantly improved progression-free survival of MM patients. However, most individuals will eventually experience disease relapse (1). The success of immunotherapy modalities greatly relies on a functionally intact patient immune system. For example, immunomodulatory imide drugs (IMiDs) can induce production of IL-2 and IFNγ by T cells, which in turn augment natural killer (NK) cell-mediated killing of myeloma cells (3). The monoclonal antibody daratumumab, targeting CD38 on myeloma cells, leads to cell death by inducing NK-cell mediated antibody-dependent cellular cytotoxicity (ADCC) (4). Furthermore, bispecific T cell engagers (BiTE) directed against B cell maturation antigen (BCMA) expressed on myeloma cells, activate T cells to eliminate tumor cells (5). Combined, these novel drugs highlight the importance of a detailed understanding of the BM immune responses to MM cells to allow for future optimization of immunological therapies for patients.

*Ex vivo* studies using human BM aspirates and biopsies have provided valuable insights into the BM microenvironment and the MM niche (6-9). However, the fact that patients upon diagnosis have systemic disease with involvement of multiple bone marrow locations makes studying the early immune events following MM cell appearance very challenging. The Radl/5T cell transfer models (10) are a series of well-established murine myeloma models used to investigate myeloma pathogenesis, response to therapy, and interaction with the BM microenvironment (11). In these models, murine 5T myeloma cells, that were derived from a spontaneously arising myeloma in a C57Bl/KaLwRij mouse, are transferred into syngeneic recipients that rapidly and reproducibly develop myeloma within 2-4 weeks. Mice present with symptoms recapitulating human disease, including tumor growth within the BM causing consequential osteolytic bone disease, presence of serum paraprotein, and renal dysfunction (12). Of the several 5T subclones that have evolved from the original clone, only the 5TGM1 cell line can be cultured *in vitro* (13). These cells have also been engineered to express eGFP for easier visualization and cell tracing (14), and are regularly used for myeloma-related studies.

Intriguingly, even though myeloma-prone C57Bl/KaLwRij mice are derived from interbreeding of C57Bl/6 mice, this HLA-identical parental strain is resistant to development of myeloma upon transfer of 5TGM1 cells (15). Since disease course of most cancers is profoundly influenced by the local tissue immune microenvironment (16), we hypothesized that alterations in BM immune status and immunological responses to MM cells might contribute to differences observed in myeloma susceptibility between C57Bl/6 and KaLwRij mice.

In this study we analyzed anti-myeloma immunity in C57Bl/6 and KaLwRij mice. Reduced incidence and severity of myeloma in C57Bl/6 mice was associated with increased cytotoxic immune responses and activation of IFNγ signaling. The latter has a central role in the 5TGM1 pathobiology as neutralization of IFNγ increases both incidence and severity of the disease. Together our data identify IFN signaling as an essential part of anti-myeloma immunity in the 5TGM1 model, and highlight the importance of the C57Bl/6 model to study multiple myeloma in an immune-competent environment.

## Materials and Methods

### Mice

8-week-old female C57Bl/6JOlaHsd and C57Bl/KaLwRijHsd mice were purchased from Envigo, The Netherlands. 5TGM1-GFP cells were kindly provided by Prof. Babatunde Oyajobi (UT Health, San Antonio, Texas, USA) (14) and were cultured in RPMI 1640 supplemented with 10% FCS, 1% PS, 1% Glutamax, 1% NEAA, and 1% Na-Pyruvate. To induce myeloma, 10^6^ 5TGM1-GFP cells were injected into the lateral tail vein of mice. Mice were sacrificed on day 21 unless stated otherwise in figure legends. Mice were euthanized when any of the following symptoms appeared: high IgG_2b_ levels, sudden and/or large weight loss, hind limb paralysis, and poor general condition; in compliance with local animal welfare guidelines. Throughout experiments mice were housed in the animal facilities of the Erasmus MC, Rotterdam, The Netherlands. Animal studies were approved by the institutional review board of the Erasmus University Medical Center as part of the animal experiment licence# AVD1010020197844.

### IFNγ neutralization

C57Bl/6JOlaHsd mice received 500 μg anti-IFNγ antibody (clone XMG1.2, rat IgG1) or rat IgG1 isotype control (clone GL113) intraperitoneally (ip) on days 0 (prior to cell transfer), 3, 7, 10, 14, and 17. Mice were sacrificed on day 21.

### M-protein measurements

50 μl of blood was collected weekly (day 0, day 7, day 14, and day 21) by cheek puncture into tubes containing K2 EDTA (Sarstedt, 16.444). ELISA was performed using a purified goat anti-mouse IgG_2b_ (A90-109A, Bethyl Laboratories, Inc) as capture antibody and an HRP-labeled goat anti-mouse IgG_2b_ (A90-109P, Bethyl Laboratories, Inc) as detection antibody (both previously sold as part of the Mouse IgG_2b_ ELISA Quantitation Set, Cat# E90-109, Bethyl Laboratories, Inc), and mouse IgG_2b_ Southern Biotech, Cat# 0104-01) as standard. Tetramethylbenzidine (TMB, Thermo Fisher) was used as substrate solution. The ELISA color reaction was measured with the Perkin Elmer VICTOR X4 Multilabel Plate reader, according to standard protocols.

### Antibodies and reagents

The following antibodies were used: CD3e-PerCpCy5.5 (eBioscience, clone 145-2C11), CD3e-PE/Dazzle594 (Biolegend, clone 145-2C11), CD8-FITC (BD, clone 53-6.7), CD11b-APCeFluor780 (eBioscience, clone M1/70), CD19-AF700 (eBioscience, clone eBio1D3), CD27-BV510 (Biolegend, clone LG.3A10), CD31-PE/CF594 (BD, clone MEC13.3), CD44-BV510 (Biolegend, clone IM7), CD45-APC/Cy7 (eBioscience, clone 30-F11), CD51-PE (eBioscience, clone RMY-7), CD69-AF647 (Biolegend, clone H1.2F3), Biotinylated CD71 (Biolegend, clone R17217), PD1-PerCP/Cy5.5 (Biolegend, clone 29F.1A12), NKp46-PE-Cy7 (eBioscience, clone 29A1.4), Tim3-PE/Cy7 (Biolegend, clone RMT3-23), Gr1-APC (BD, clone RB6-8C5), Ly6c-BV510 (Biolegend, clone HK1.4), Lsel-AF700 (Biolegend, clone MEL-14), Sca-1-PerCP/Cy5.5 (Invitrogen, clone D7), Ter119-BV510 (Biolegend, clone TER-119), Streptavidin-PE/Cy7 (Biolegend).

### Flow cytometry and fluorescence activated cell sorting

For fluorescence activated cell sorting (FACS) of bone marrow stromal cells two femurs and two tibias from each mouse were used. The ends of cleaned bones were cut off and marrow was flushed out. The remaining bone was cut into small pieces and gently crushed with a mortar and pestle. Bone fragments were filtered through a 100 μm cell strainer and then digested for 45 minutes at 37C with 0.25% Collagenase Type I (Stem Cell). Bone chips were again washed on a 100 μm cell strainer and stromal cells were collected into PBS containing 1% FCS and 1 mM EDTA. Erythrocytes were lysed and cells were counted. For flow cytometry and FACS of bone marrow immune cells 1 femur and 2 tibias from each mouse were used. After isolating and cleaning the bones, one end was cut off and bones were placed into a 0.5 ml microcentrifuge tube with a hole punched in the bottom. This was nested into a 1.5 ml microcentrifuge tube and spun down at 10,000 RPM for 15 seconds to collect the bone marrow (11). Following resuspension in PBS containing 1% FCS and 1 mM EDTA, erythrocytes were lysed. Remaining cells were washed with PBS containing 2% FCS, filtered through a 100 μm cell strainer and counted. Spleens were mechanically disrupted to obtain a single cell suspension. Following erythrocyte lysis, the samples were filtered through a 100 μm cell strainer and counted. For all flow cytometry and FACS experiments, Fc-receptors were blocked with 10% normal mouse serum and anti-CD16/32 (BD, clone 2.4G2) prior to incubation with appropriate antibodies for 25 minutes on ice in the dark. Following staining, cells were washed and resuspended in PBS + 2% FCS containing DAPI, and measured on a BD LSRII or sorted with a BD FACS Aria III. Results were analyzed using FlowJo v10.

### Single-cell RNA sequencing

Sorted T (CD3^+^) and B (CD19^+^) cells, NK cells (CD3^-^CD19^-^NKp46^+^), monocytes/macrophages (CD3^-^CD19^-^NKp46^-^CD11b^+^Gr1^lo^LyC6^int/hi^) and neutrophils (CD3^-^CD19^-^NKp46^-^ CD11b^+^Gr1^+^LyC6^int^) were counted using a Countess II Automated Cell Counter (Thermo Fisher) and then combined and processed as one sample per mouse. Single cells were encapsulated for cDNA synthesis and barcoded using the Chromium Single Cell 3’ Reagent Kit v3 (10X Genomics), followed by library construction according to the manufacturer’s recommendations. The Agilent 2100 Bio-Analyzer and the High Sensitivity DNA kit was used to determine the quality and quantity of libraries, which were then sequenced on the NovaSeq 6000 platform (Illumina), paired-end mode, at a sequencing depth of ∼20,000 reads/cell, followed by computational alignment using CellRanger (v3.0.2, v4.0.0, and v6.0.1, 10x Genomics). Datasets were subjected to quality control steps using Seurat (R package, v 4.0) (17), including selecting cells with a library complexity of more than 200 features, removing doublets, and filtering out cells with high percentages of mitochondrial genes (>10%). NK cells (*Ncr1+*) and T cells (*Cd3e+*) were subsetted from each mouse using the subset() function, and merged by integration and label transfer (18) into one NK/T dataset, and normalized and analyzed by principal-component analysis (PCA) on the most variable genes (k = 2,000) across all cells to implement dimensionality reduction. Samples were integrated in the following order: first the biological duplicates, then the two different conditions (uninjected vs tumor-bearing) per strain, and finally the two strains. From this dataset, NK cells were subsetted and unsupervised clustering was performed using a shared nearest neighbor (SNN) modularity optimization-based clustering algorithm (resolution 0.3-0.4) (19). Cells were projected in two dimensions using Uniform Manifold Approximation and Projection (UMAP) (20). Significantly upregulated genes in each cluster compared to all other clusters (adjusted p <0.05) were identified using the Seurat Function FindMarkers.

### Bulk RNA sequencing

RNA was isolated from sorted Nkp46^+^ or CD8^+^ cells with the NucleoSpin RNA XS kit (Macherey-Nagel) and cDNA was prepared using the SMARTer Ultra Low RNA kit (Clontech Laboratories) for Illumina Sequencing following the manufacturer’s protocol. The Agilent 2100 Bio-Analyzer and the High Sensitivity DNA kit was used to verify quality and quantity of cDNA samples. cDNA libraries were generated using the TruSeq Sample Preparation v.2 Guide (Illumina) and paired end-sequenced on the NovaSeq 6000 (Illumina). Adaptor sequences and polyT tails were trimmed from unprocessed reads using fqtrim version v0.9.7 (http://ccb.jhu.edu/software/fqtrim/). Read counts and TPMs were determined with Salmon version v1.4.0 (21). Per gene pseudocount estimates were determined from the Salmon output with the tximport R package (22). These per gene pseudocount estimates were in turn used for determining differential expression of genes between groups of interest through the DESeq2 package (23). GSEA was performed with the GSEA software (Broad Institute) using predefined gene sets from the Molecular Signatures Database (MSigDB 7.2). Gene lists were ranked on the basis of the shrunken log 2-fold change based on the ashr method (24) made available through the DESeq2 package. Classical enrichment statistics with 10000 permutations was used to determine significant enrichment within gene sets. Gene Ontology analysis was done with the PANTER Overrepresentation Test (GO database release: 2019-09-03).

### Statistics

Statistical analyses were performed in GraphPad Prism9 or in R v4.1. Statistical significance was assessed using Mann-Whitney *U* tests, Fisher’s exact test, Pearson’s test, or Wald test followed by a Benjamini-Hochberg correction. ns, p>0.05; *, p<0.05; **, p<0.01; ***, p<0.001; ****, p<0.0001.

### Data availability

The single-cell RNA sequencing datasets of bone marrow NK cells from control and tumor-bearing C57Bl/6JOlaHsd and C57Bl/KaLwRijHsd mice are available at ArrayExpress, no. E-MTAB-12616. This dataset is also available for interactive visualization at www.bmbrowser.org. The bulk RNA sequencing datasets of bone marrow NK cells, CD8 T cells, and stromal cells are available at ArrayExpress, no. E-MTAB-12615. The scripts generated during this study are available through GitHub at https://github.com/MyelomaRotterdam/Kellermayer-et-al-2023.

## Results

### Development of myeloma in C57Bl/6 mice

To analyze MM development in C57Bl/6 mice, GFP-labeled 5TGM1 cells were intravenously injected and peripheral blood was analyzed weekly to determine IgG_2b_ M-protein level as a measure for tumor growth (**Fig1A**). As a positive control, KaLwRij mice were used as recipients. As previously reported (13), incidence of myeloma in KaLwRij mice was 100%, with elevated levels of IgG_2b_ from day 14 post-inoculation onwards (median at day 21: 4.54 mg/ml) (**Fig1B**). In contrast, at day 14, C67Bl/6 mice showed no signs of myeloma development and IgG_2b_ levels were similar to baseline (**Fig1B**). However, at day 21, IgG_2b_ levels in C67Bl/6 mice were elevated and comparable to the levels in KaLwRij mice at day 14 (**Fig1B**). Experiments were terminated at day 21, the time when KaLwRij mice reached humane endpoints. At this time point, none of the C67Bl/6 mice had reached humane endpoints. In line with the elevated M-protein levels, a subset of the C57Bl/6 mice contained scattered aggregates of GFP^+^ tumor cells in femurs, while KaLwRij mice had significant numbers of GFP^+^ cells (**Fig1C**). Flow cytometric assessment of BM cells showed that 59% (13 out of 22) of C57Bl/6 mice developed myeloma, defined as >0.01% GFP^+^ tumor cells of all live cells in the flushed BM samples (**Fig1D-E**). The percentage of BM GFP^+^ cells in C57Bl/6 mice was significantly lower compared to KaLwRij (**Fig 1E**), also when including only tumor-bearing C57Bl/6 mice (**Supplementary Fig1A**). In addition, two distinct tumor growth patterns were appearent. The vast majority of KaLwRij mice (85%, 18 out of 21 mice) presented with unrestrained BM growth, defined as >5% GFP^+^ cells in the BM, while the remainder of the KaLwRij animals showed restrained BM growth with GFP percentages between 0.01 – 5% of live BM cells (**Fig1E-F**). Conversely, in tumor-bearing C57Bl/6 mice, the majority of mice had a restrained tumor (0.01 – 5% GFP of live BM cells), and only 23% (3 out of 13) of animals presented with the unrestrained growth pattern (**Fig1E-F, Supplementary Table I**).

**Figure 1.**
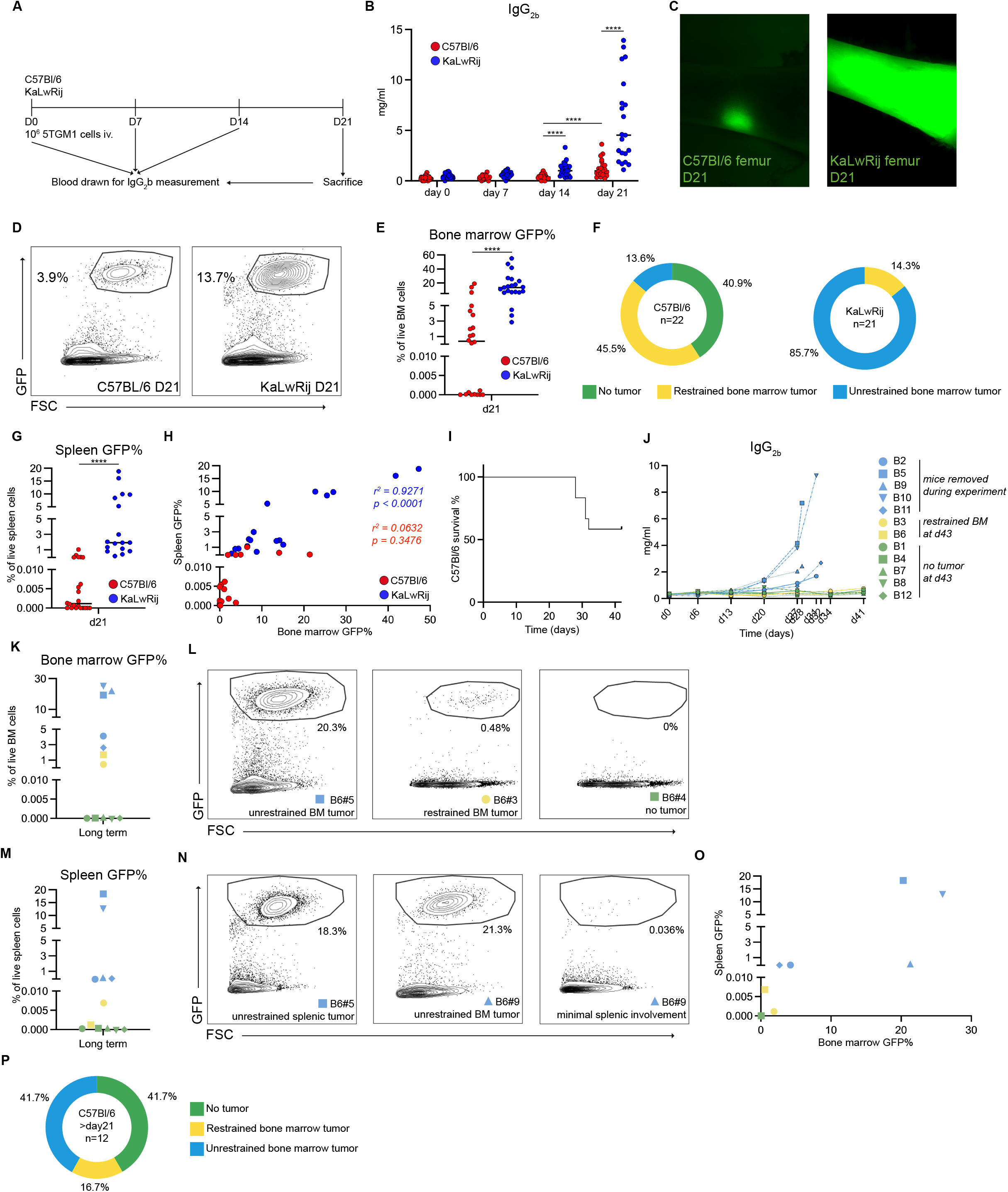
Multiple myeloma development in C57Bl/6 compared to KaLwRij mice. **A**, Experimental setup of the 5TGM1 transfer model. D: day; iv: intravenous injection. **B**, IgG2b levels in serum after 5TGM1 transfer. **C**, Representative images of GFP in C57Bl/6 and KaLwRij femurs 21 days after injection of tumor cells, 5x magnification. **D**, Representative flow cytometric plots of bone marrow from C57Bl/6 and KaLwRij. **E**, Percentage of GFP+ tumor cells in bone marrow at day 21. **F**, Donut graphs depicting myeloma growth patterns in C57Bl/6 (n=22) and KaLwRij (n=21) mice. Green, no tumor; yellow, restrained bone marrow tumor; blue, unrestrained bone marrow tumor. **G**, Percentage of spleen GFP+ tumor cells at day 21. **H**, Correlation between bone marrow and spleen tumor percentage. Red, r2 value for C57Bl/6; blue, r2 value for KaLwRij. **I**, Survival curve of C57Bl/6 mice post day 21. **J**, IgG2b levels from C57Bl/6 mice allowed to progress beyond day 21. Blue, mice removed during the experiment; yellow, mice with restrained bone marrow tumor at day 43; green, mice with no tumor at day 43. **K**, Percentage of bone marrow GFP+ tumor cells in individual C57Bl/6 mice allowed to progress beyond day 21. **L**, Representative flow cytometric plots showing bone marrow from mice with unrestrained bone marrow tumor (B6#5, left), restrained bone marrow tumor (B6#3, middle), or no tumor (B6#4, right). **M**, Percentage of spleen GFP+ tumor cells in individual C57Bl/6 allowed to progress beyond day 21. **N**, Representative flow cytometric plots showing spleen from a mouse with high splenic involvement (B6#5, left), bone marrow from a mouse with unrestrained bone marrow involvement (B6#9, middle) and spleen from the same mouse (B6#9, right) presenting with low splenic involvement. **O**, Correlation between bone marrow and spleen tumor percentage in C57Bl/6 allowed to progress beyond day 21. **P**, Donut graph depicting myeloma growth patterns in C57Bl/6 allowed to progress beyond day 21. Green, no tumor; yellow, restrained bone marrow tumor; blue, unrestrained bone marrow tumor. Data in A-H are from at least 3 independent experiments. Line indicates median of values. Statistical significance was assessed using Mann-Whitney *U* tests (ns, *p*>0.05; *, *p*<0.05; **, *p*<0.01; ***, *p*<0.001; ****, *p*<0.0001).

Even though myeloma cells are rarely found in the spleen of patients, splenic dissemination is a well-known feature of 5TGM1 cells when injected into KaLwRij mice (25). Accordingly, all KaLwRij mice (n = 21) had splenic involvement, defined as GFP^+^ cells >0.01% of all live spleen cells (**Fig1G**), and accompanied by a significant increase in spleen weight (**Supplementary Fig1B**). In contrast, half of the tumor-bearing C57Bl/6 mice (53.8%, 7 out of 13) had no splenic involvement (**Supplementary Table I**), and in the C57Bl/6 mice with splenic involvement the splenic tumor load (defined as % of GFP^+^ cells) was significantly lower compared to KaLwRij mice (**Supplementary Fig1C**). In KaLwRij mice splenic tumor involvement showed a significant positive correlation with the BM tumor load (r^2^ = 0.9271, p < 0.0001) while this was not the case in C57Bl/6 mice (r^2^ = 0.0632, p = 0.3476) (**Fig1H**).

Based on IgG_2b_ levels, kinetics of tumor development in C57Bl/6 mice lagged by approximately 1 week compared to KaLwRij mice (**Fig1B**). This prompted us to extend the period during which 5TGM1 tumors were allowed to grow in C57Bl/6 mice. From day 21 onwards, mice were visually inspected two times per day and upon appearance of a previously defined set of symptoms (described in Materials and Methods), animals were euthanized. Between day 28 and day 32, five out of 12 C57Bl/6 mice (41.7%) developed unrestrained tumor growth (**Fig1I**). These animals had a median of 2.6 mg/ml plasma IgG_2b_ (**Fig1J**, euthanized mice indicated with blue symbols) and a median of 20.3% GFP cells in the BM with three mice having values above 20% (**Fig1K-L**). In addition, all 5 mice had developed splenic involvement based on our previous criteria (set as splenic GFP% > 0.01%) (**Fig1M**). Similar to day 21, there was no strong correlation between BM and spleen GFP content (**Fig1N-O**).

The remaining seven mice did not develop signs of distress up until 6 weeks post injection, when the experiment was concluded. Subsequent flow cytometric analyses indicated that 2 of the seven remaining mice had a restrained BM tumor (>0.01 - 5% GFP) with splenic involvement being below the threshold of 0.01%. The remaining 5 mice had no detectable tumor cells (**Fig1J-O**, mice with restrained BM tumor indicated with green symbols; mice with no tumor indicated with yellow symbols). Altogether, after 6 weeks 58.3% of C57Bl/6 mice (7 out of 12) had developed a tumor, indicating that incidence at 6 weeks was similar to incidence at 3 weeks (**Fig1P, Supplementary Table I**). However, with time, tumor burden seemed to increase, with a trend towards more mice developing an unrestrained BM tumor (71.4%, 5 out of 7 mice vs 23.1%, p=0.0623) at the expense of mice with a restrained tumor (28.6%, 2 out of 7 mice vs 76.9%).

In sum, these data indicate that approximately 60% of C57Bl/6 mice develop myeloma after intravenous transfer of 10^6^ GFP-labeled 5TGM1 cells and that the severity of disease in tumor-bearing mice increases with time, suggesting the possibility of initial immune-mediated tumor control that gets overwhelmed as the tumor grows.

### scRNAseq analysis of NK cells in myeloma-bearing mice

We hypothesized that delayed tumor kinetics in C57Bl/6 mice were due to immune-mediated containment. Therefore, we set out to define anti-myeloma immune responses by single-cell RNA-sequencing (scRNA-seq). Single-cell RNA libraries were generated from a pool of purified BM NK and T cells (together with B cells, neutrophils and monocytes for future analyses, sorting strategy described in **Supplementary Fig2A**) from non-injected C57Bl/6 and KaLwRij mice and from mice 21 days after iv injection of 5TGM1 cells that had developed a restrained (C57Bl/6) or unrestrained (KaLwRij) tumor. Characteristics of the mice used for scRNA-seq are in **Supplementary Table II**. After quality control and preprocessing, a total number of 45,623 transcriptomes from 8 mice at two time points were obtained (**Supplementary Fig2B**). To increase resolution, we identified and subsetted NK (*Ncr1*) and T (*Cd3e*) cells *in silico* and integrated the objects of the respective cell types for downstream analyses (**Supplementary Fig2C**).

The NK cell object contained 5,126 cells in 10 clusters (**Fig2A**), including small numbers of contaminating neutrophils (*Ngp, Retnlg*) and B cells (*Ms4a1, Cd19*), which were excluded from downstream analyses (**Fig2B**). Recently, continuous stages of NK cell development stages have been transcriptomically characterized in murine bone marrow (26). Based on this published transcriptomic dataset, we identified proliferating NK cells (transcribing high levels of *Mki67* and *Stmn1*), transitional NK1 cells (*Eif5a, Mif, Nme1*), transitional NK2 cells (*Klra4, Klra8*), transitional NK3 cells (*Ccl3, Xcl1, Nr4a1*), terminally differentiated NK cells (*Irf8, Lgals1, Cma1*), and interferon-responsive NK cells (*Isg15, Ifit3*) (**Fig2B**). Besides NK cells we also observed a cluster of *Tmem176a-* and *Tmem176b-*expressing ILCs (‘Tmem176^+^ ILC’) and another cluster that showed low transcription of *Tmem176a* and *Tmem176b* but nevertheless transcribed other ILC1-related genes (*Il7r, Cxcr6, Ikzf2;* ‘ILC1’) (27).

Based on surface markers, NK follow a maturation path starting at CD11b^low^CD27^low^ cells, followed by upregulation of CD27 (CD11b^low^CD27^high)^ and upregulation of CD11b (CD11b^high^CD27^high^) before losing CD27 in the most mature cells (CD11b^high^CD27^low^) (28). In this scheme, CD11b expression serves as a marker of functional maturity (29). Based on gene expression levels of *Itgam* (encoding CD11b), *Cd27, Klrg1*, and *S1pr5* (the latter two expressed on terminally differentiated NK cells (30,31)) (**Supplementary Fig3A**), we labeled the ‘Transitional NK1’ and ‘Transitional NK3’ clusters as immature NK cells (both being Cd11b-), and the ‘Transitional NK2’ (Cd11b^+^Cd27^+^) and ‘Terminally differentiated’ (CD11b^high^CD27^low^) clusters as mature NK cells, respectively (**Supplementary Fig3B**). Split UMAP representations (**Fig2C**) and enumerations (**Fig2D-E, Supplementary Fig 3C**) revealed that at steady state (SS) the frequency of total mature NK cells was comparable between the two strains. Upon tumor inoculation, the percentage of mature NK cells increased in tumor-bearing C57Bl/6 mice compared to steady state (**Fig2E, Supplementary Fig3D**). In sharp contrast, the frequency of these mature cells declined in KaLwRij mice upon tumor transfer. Interestingly, this decrease was coupled with a robust increase of the ‘ILC1’ cluster in KaLwRij, but not in C57Bl/6, bone marrow.

**Figure 2.**
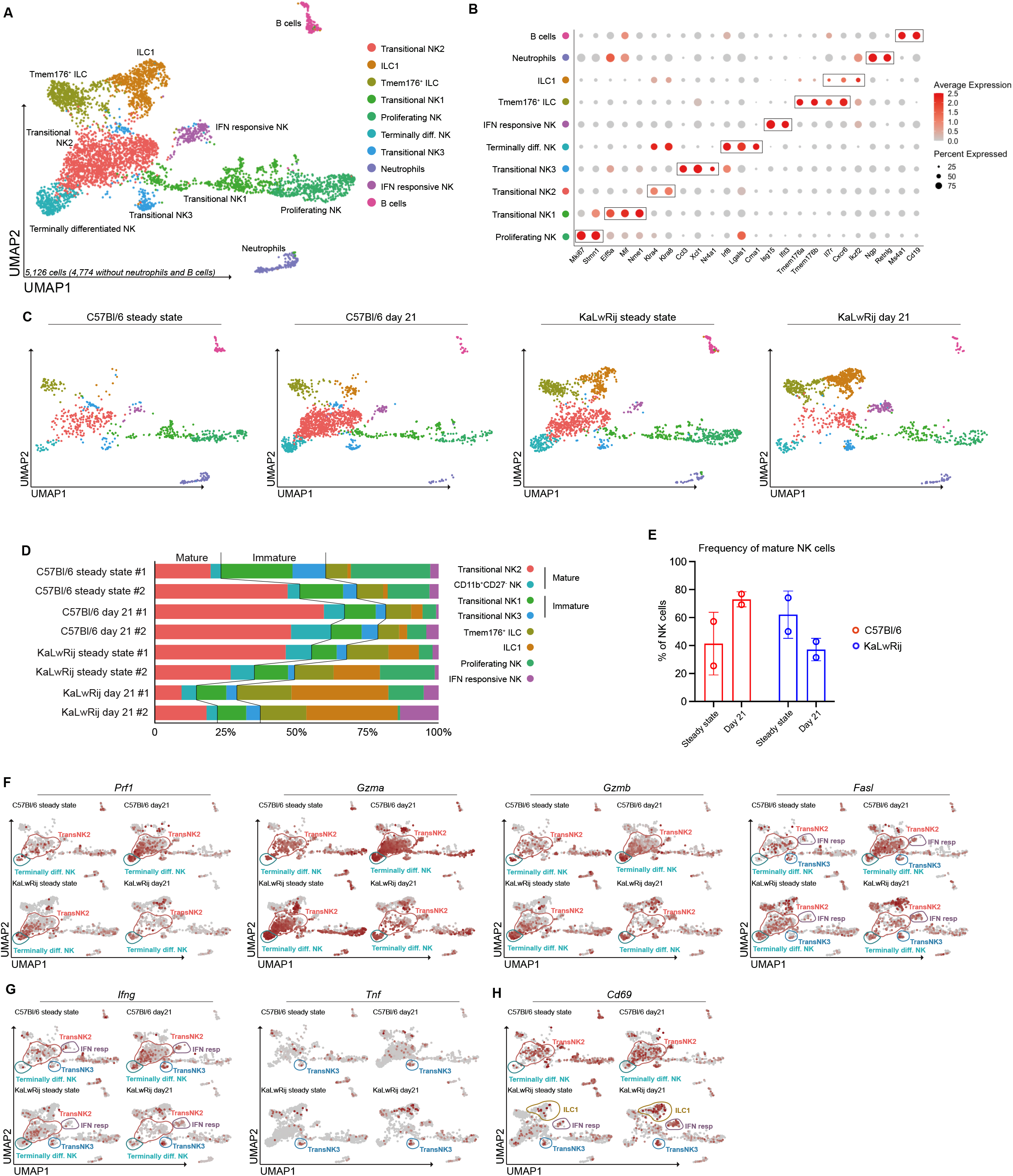
Single-cell RNAseq of NK cells in C57Bl/6 and KaLwRij mice after 5TGM1. **A**, UMAP of NK cells from C57Bl/6 and KaLwRij mice. Two animals per strain at steady state and 2 animals per strain at d21 after 5TGM1. Colors indicate clusters obtained at resolution 0.4. **B**, Dot plot depicting average expression levels of canonical genes used to identify clusters. **C**, Split UMAP of NK cells per condition. **D**, Bar graph depicting cluster distribution in individual mice. Colors indicate clusters. **E**, Bar graphs showing frequencies of grouped mature clusters under steady state condition and day 21 after tumor transfer. **F**, Feature plots split per condition of genes associated with NK cell cytotoxicity; circled areas refer to indicated clusters colored using respective UMAP colors. **G**, Feature plots split per condition of cytokines associated with NK activity; circled areas refer to indicated clusters colored using respective UMAP colors. **H**, Feature plot split per condition of Cd69 activation marker; circled areas refer to indicated clusters colored using respective UMAP colors.

We next set out to characterize functional features of the NK cell transcriptome. NK cells have been grouped into cytotoxic NK cells and cytokine producing NK cells (32). The frequency of cells transcribing genes encoding the cytotoxic proteins perforin (encoded by *Prf1*), granzyme A (encoded by *Gzma*), granzyme B (encoded by *Gzmb*) and *Fasl* (**Fig2F**) increased in tumor-bearing C57Bl/6 mice after tumor inoculation, but decreased in KaLwRij mice (**Fig2F**). Cytokine-producing NK cells mainly secrete TNF*α* and IFNγ and for both cytokines, the amount of positive cells showed a slight increase in C57Bl/6 mice upon tumor transfer, with minimal change observed in KaLwRij mice (**Fig2G**). Interestingly, the activation marker *Cd69* was transcribed by mature ‘Transitional NK2’ and ‘Terminally differentiated NK’ clusters in C57Bl/6 mice, while in KaLwRij mice *Cd69* transcription was largely absent from these clusters, yet transcribed in the ‘IFN-responsive NK’, ‘Transitional NK3’, and ‘ILC1’ cluster (**Fig2H**), suggestive of differential NK cell/ILC activation.

Besides their preferential activation in KaLwRij mice after tumor inoculation, ILC1 cluster were also numerically overrepresented in KaLwRij mice at steady state, and their abundance increased even further upon tumor development (**Fig2C-D**). Although this cluster was characterized by genes associated with ILC1s (*Ikzf2* [encoding Helios], *Cxcr6*, and *Il7r)* (27) (**Fig2B**), feature plots for these genes suggested heterogeneity within the cluster (**Supplementary Fig3E**). Therefore, we subsetted and reanalyzed this cluster (**Supplementary Fig3F**). This indeed revealed a cluster transcribing ILC1-related genes, consisting mainly of cells with KaLwRij origin; but also a second cluster resembling the ‘Transitional NK2’ cluster from the original dataset (named ‘Transitional NK2-like’) (**Supplementary Fig3G**). This ‘Transitional NK2-like’ cluster was characterized by transcription of cytotoxicity-related genes and contained cells from both mouse strains (**Supplementary Fig3G-H**). Split UMAPs and enumeration of cluster contribution to the total NK cell pool revealed a clear overrepresentation of the ‘ILC1’ cluster in KaLwRij mice compared to C57Bl/6 (**Supplementary Fig3H-I**).

Together these transcriptomic data identify differential NK cell and ILC1 distribution and activation following tumor inoculation in C57Bl/6 and KaLwRij mice.

For T cells, there was a large variation in the number of both sorted and *in silico* recovered cells per mouse, in part due to the combined sorting of T and B cells. This precluded any reliable analyses of the T cell compartment (**Supplementary Table II**).

### Myeloma progression is associated with loss of activated NK cells

As our transcriptomic analyses suggested differential NK cell responses in the two mouse strains, we next analyzed BM immune response to transferred tumor cells by flow cytometry. At steady state, total BM cellularity did not differ between C57Bl/6 and KaLwRij mice (**Fig3A**), while in both strains, unrestrained BM myeloma growth at day 21 led to significantly decreased numbers of non-tumor BM cells. At steady state, NK cell (NKp46^+^CD3^-^CD19^-^ cells) frequencies, but not absolute numbers, were significantly higher in C57Bl/6 mice (**Fig3B**), and this remained stable in animals with restrained tumor growth. In contrast, in KaLwRij mice at day 21, relative and absolute numbers of NK cells were significantly reduced compared to steady state. Loss of NK cells was associated with unrestrained tumor growth, as both relative and absolute NK cell numbers in C57Bl/6 mice with unrestrained tumor growth decreased to levels similar to those in KaLwRij mice (**Fig3B**).

**Figure 3.**
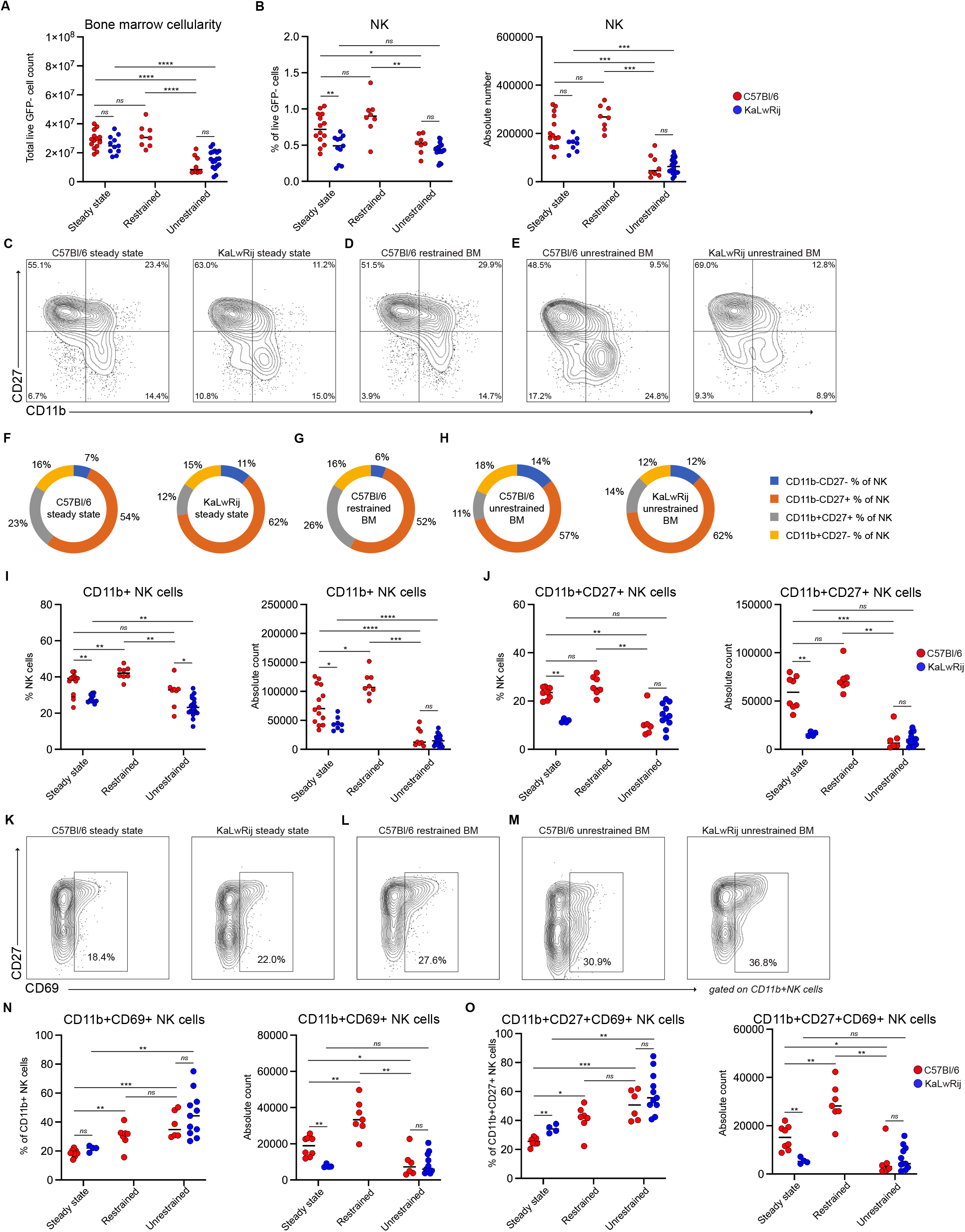
Unrestrained bone marrow growth is associated with reduction of activated NK cells. **A**, Absolute number of bone marrow live GFP-cells. **B**, Relative (left) and absolute (right) numbers of bone marrow NK cells. **C-E**, Representative flow cytometric contour plots showing bone marrow NK cell maturation stages in indicated conditions. **F-H**, Donut charts depicting bone marrow NK cell composition in indicated conditions. **I**, Relative (left) and absolute (right) numbers of bone marrow CD11b+ NK cells. **J**, Relative (left) and absolute (right) numbers of bone marrow CD11b+CD27+ NK cells. **K-M**, Representative flow cytometry contour plots showing bone marrow CD11b+ NK cell activation status in indicated conditions. **N**, Relative (left) and absolute (right) number of CD11b+CD69+ NK cells. **O**, Relative (left) and absolute (right) number of CD11b+CD27+CD69+ NK cells. Data are from at least 3 independent experiments. Line indicates median of values. Statistical significance was assessed using Mann-Whitney *U* tests (ns, *p*>0.05; *, *p*<0.05; **, *p*<0.01; ***, *p*<0.001; ****, *p*<0.0001).

To identify NK cell populations differentially present at steady state we used CD11b and CD27 to define the major developmental NK subsets (**Fig3C-J**). The percentage of mature CD11b^+^ NK cells (corresponding with the mature ‘Transitional NK2’ and ‘Terminally differentiated’ NK clusters from the sc-transcriptomic dataset) were significantly higher in C57Bl/6 mice at steady state compared to KaLwRij (**Fig3C, F, I**), and this difference was due to alterations in the highly cytotoxic CD11b^high^CD27^high^ NK cell subset (29) (**Fig3J)** but not the CD11b^low^CD27^hi^ or CD11b^hi^CD27^low^ subset (**Supplementary Fig4A-B**). A similar comparison was made to identify the NK cell population lost in mice with unrestrained MM. In response to 5TGM1 MM cells, C57Bl/6 mice with restrained BM tumor had a modest increase in relative, and a pronounced increased in absolute numbers of mature CD11b^+^ NK cells (**Fig3D, G, I**). In contrast, upon development of unrestrained myeloma growth, frequency and absolute number of total CD11b^+^ and CD11b^high^CD27^high^ NK cells were significantly reduced, comparable to KaLwRij mice (**Fig3E, H, I, J**). This decrease was also significant among total GFP^-^ cells, indicating that loss of mature cytotoxic NK cells is not a general consequence of decreased cell numbers due to the increased presence of myeloma (**Supplementary Fig4C**). Inversely, the percentage of CD11b^-^ NK cells (corresponding with the immature ‘Transitional NK1’ and ‘Transitional NK3’, and also including the ‘Tmem176^+^ ILC’ and ‘ILC1’ subsets from the transcriptomic data) increased in mice with unrestrained BM tumors, and was higher in KaLwRij mice compared to C57Bl/6 mice (**Supplementary Fig4D**). However, this relative increase was not coupled with an increase in absolute numbers of CD11b^-^ NK cells.

To analyze whether the differential kinetics of NK cells were also related to the altered activation that we had observed on the transcriptome level, we determined CD69 protein expression on mature NK cell as a proxy for cellular activation (33) (**Fig3K-M, Supplementary Fig4E**). The percentages of activated total mature CD11b^+^ NK cells (**Fig3N**) and in specific activated mature CD11b^high^CD27^high^ NK cells (**Fig3O**) increased upon tumor presence in both mouse strains. In C57BL/6 mice with restrained tumor growth this increase in percentage was accompanied by an increase in absolute cell numbers (**Fig3N, O**). In contrast, unrestrained tumor growth was associated with a loss of absolute numbers of activated NK cells, despite higher percentages (**Fig3N, O, Supplementary Fig4F-G**). A similar disparity between percentage and absolute numbers of activated NK cells was seen in KaLwRij mice, again indicative of a correlation with unrestrained tumor outgrowth. The changes in relative and absolute numbers of CD11b^-^CD69^+^ NK cells were similar to that observed in the CD11b^+^ NK cells (**Supplementary Fig4H**).

### Myeloma progression is associated with loss of activated CD8 T cells

We next investigated whether similar associations between cellular activation and tumor behavior were apparent for cytotoxic T cells. At steady state, both the percentage and absolute number of BM CD8^+^ T cells were higher in C57BL/6 mice compared to KaLwRij mice (**Fig4A**). During restrained tumor growth in C57BL/6 mice, relative and absolute numbers of CD8 T cells remained constant, with a slight increase in absolute numbers explained by changes in BM cellularity as a result of tumor presence (**Fig3A, Supplementary Fig5A-B**). Unrestrained tumor growth in C57Bl/6 mice was accompanied by a significant decrease in the absolute numbers of CD8 T cells (**Fig4A**). This was similar to KaLwRij mice, in which numbers and percentages of CD8 T cells during unrestrained tumor growth decreased below the already lower steady state levels (**Fig4A**).

**Figure 4.**
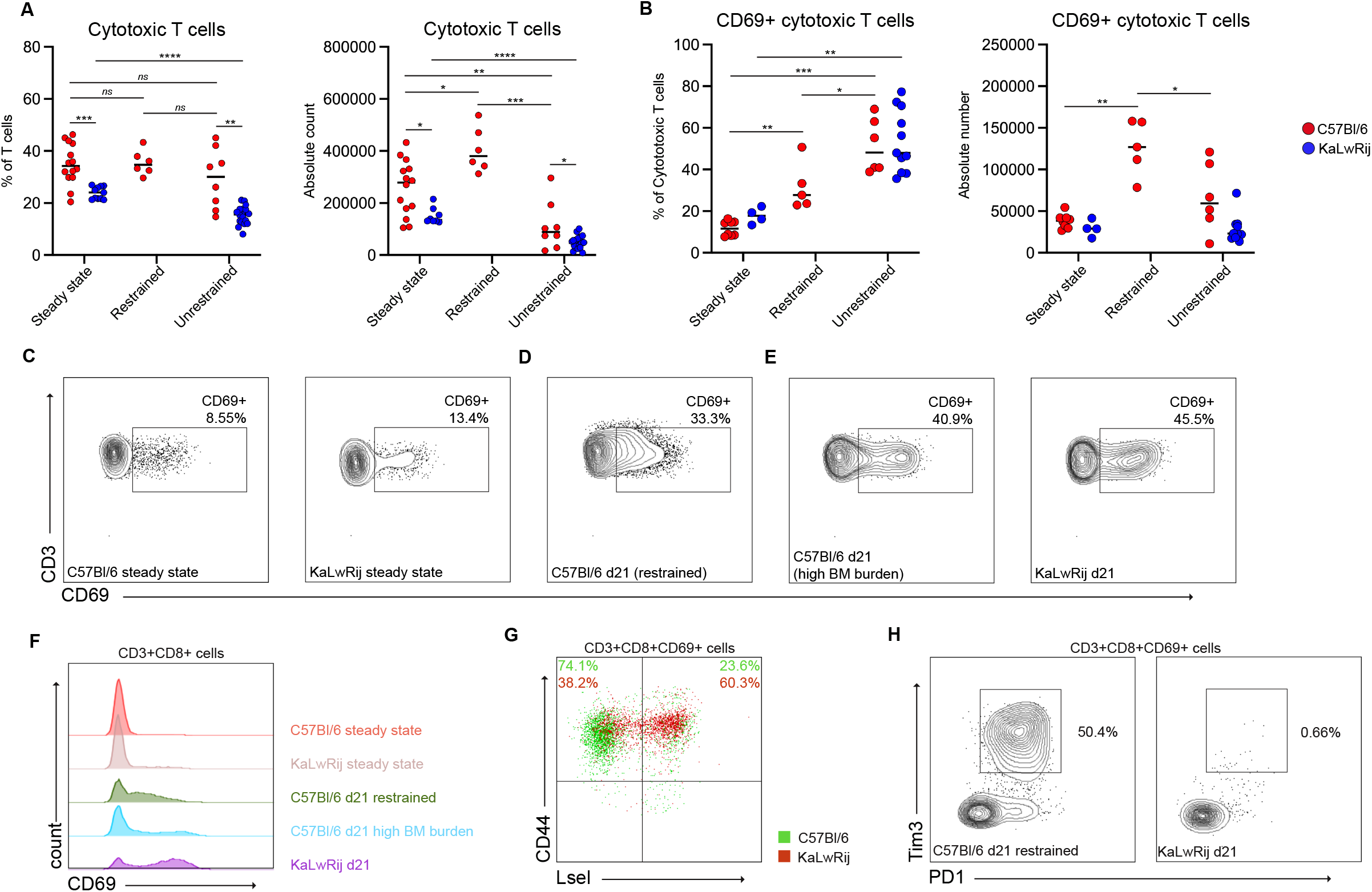
Unrestrained bone marrow growth is associated with reduction of activated CD8 T cells. **A**, Relative (left) and absolute (right) number of cytotoxic T cells. **B**, Relative (left) and absolute (right) number of CD69+ cytotoxic T cells. **C-E**, Representative flow cytometry plots showing activation status of cytotoxic T cells in indicated conditions. **F**, Flow cytometry histograms from representative samples showing CD69 expression on cytotoxic T cells. **G**, Representative flow cytometry plot showing CD44/L-Selectin status of bone marrow CD69+ cytotoxic T cells from C57Bl/6 (green) and KaLwRij (red) mice. **H**, Representative flow cytometry contour plots showing Tim3/PD1 status of bone marrow CD69+ cytotoxic T cells from C57Bl/6 (left) and KaLwRij (right) mice. Data in A-F are from at least 3 independent experiments; data in G-H are representative stainings from at least 2 independent experiments. Line indicates median of values. Statistical significance was assessed using Mann-Whitney *U* tests (ns, *p*>0.05; *, *p*<0.05; **, *p*<0.01; ***, *p*<0.001; ****, *p*<0.0001).

To analyze activation status of BM cytotoxic T cells we determined CD69 expression. At steady state minimal numbers of CD69 expressing cytotoxic T cells were present in both strains (**Fig4B-C**). Upon tumor presence, percentage of activated CD8 T cells increased in C57BL/6 mice with restrained tumor growth, and increased even further in mice with unrestrained tumor growth, similar to KaLwRij mice (**Fig4B, D-E**). The absolute number of activated CD8 T cells increased significantly in C57Bl/6 mice with a restrained tumor, and although it did decrease in mice with an unrestrained tumor, inter-mouse variability was high. Furthermore, when measured as % of GFP^-^ cells, the frequency of CD8^+^CD69^+^ T cells in C57Bl/6 mice with an unrestrained tumor did not differ from mice with restrained tumor and remained significantly higher compared to C57Bl/6 steady state and tumor-bearing KaLwRij mice (**Supplementary Fig5C**).

Interestingly, while in restrained tumor-bearing C57Bl/6 mice CD69 expression showed a gradual increase (**Fig4D**), cytotoxic T cells from all tumor-bearing KaLwRij mice, and unrestrained C57Bl/6 mice with highest BM tumor burden, expressed only CD69 at high levels (**Fig4E-F**), suggestive of altered or incomplete activation. To further characterize BM CD69^+^ cytotoxic T cells, other commonly used activation markers were analyzed. In C57Bl/6 mice with a restrained tumor, CD69^+^ T cells mainly had an activated CD44^+^L-Selectin^−^ phenotype. In contrast, in the presence of unrestrained tumor growth in KaLwRij mice, CD69^+^ cells were CD44^+^L-Selectin^+^ (**Fig4G**). Furthermore, CD69^+^ T cells of restrained tumor marrows expressed PD1 and Tim-3 to varying degrees, while these markers were virtually absent on T cells from KaLwRij marrows (**Fig4H**). There was no association between splenic tumor burden and alterations in splenic cytotoxic immune cells (**Supplementary Fig5D-F**).

### IFNγ limits incidence and severity of 5TGM1 myeloma

Analyses of the BM immune response to 5TGM1 MM cells indicated that alterations in the cytotoxic immune response were associated with disease progression. To gain further insights into the mechanistic underpinnings of anti-myeloma immune responses we purified NK cells, cytotoxic T cells and CD45^-^CD31^-^Ter119^-^ mesenchymal stromal cells from steady state and day 21 post 5TGM1-inoculation C57Bl/6 mice with a restrained tumor (sorting strategy in **Supplementary Figs 6A and 6D)** and performed RNA-sequencing. This allowed for deeper sequencing compared to scRNAseq, and provided better coverage and detection of immune effector transcripts including cytokines. Ranked gene set enrichment analysis (GSEA) on differentially expressed genes after tumor inoculation was performed to identify pathways and genes associated with tumor responses. At day 21, NK cells were enriched in pathways related to metabolism and cell-cycle (**Fig5A**). In addition, GSEA indicated active immune effector pathways including ‘Interferon gamma response’ and ‘Interferon alpha response’ (**Fig5B-D, Supplementary Fig6B**). Analysis of cytotoxic T cells revealed activation of similar pathways, including metabolic pathways and the ‘Interferon gamma response’ and ‘Interferon alpha response’ gene sets (**Fig5E-H, Supplementary Fig6C**).

**Figure 5.**
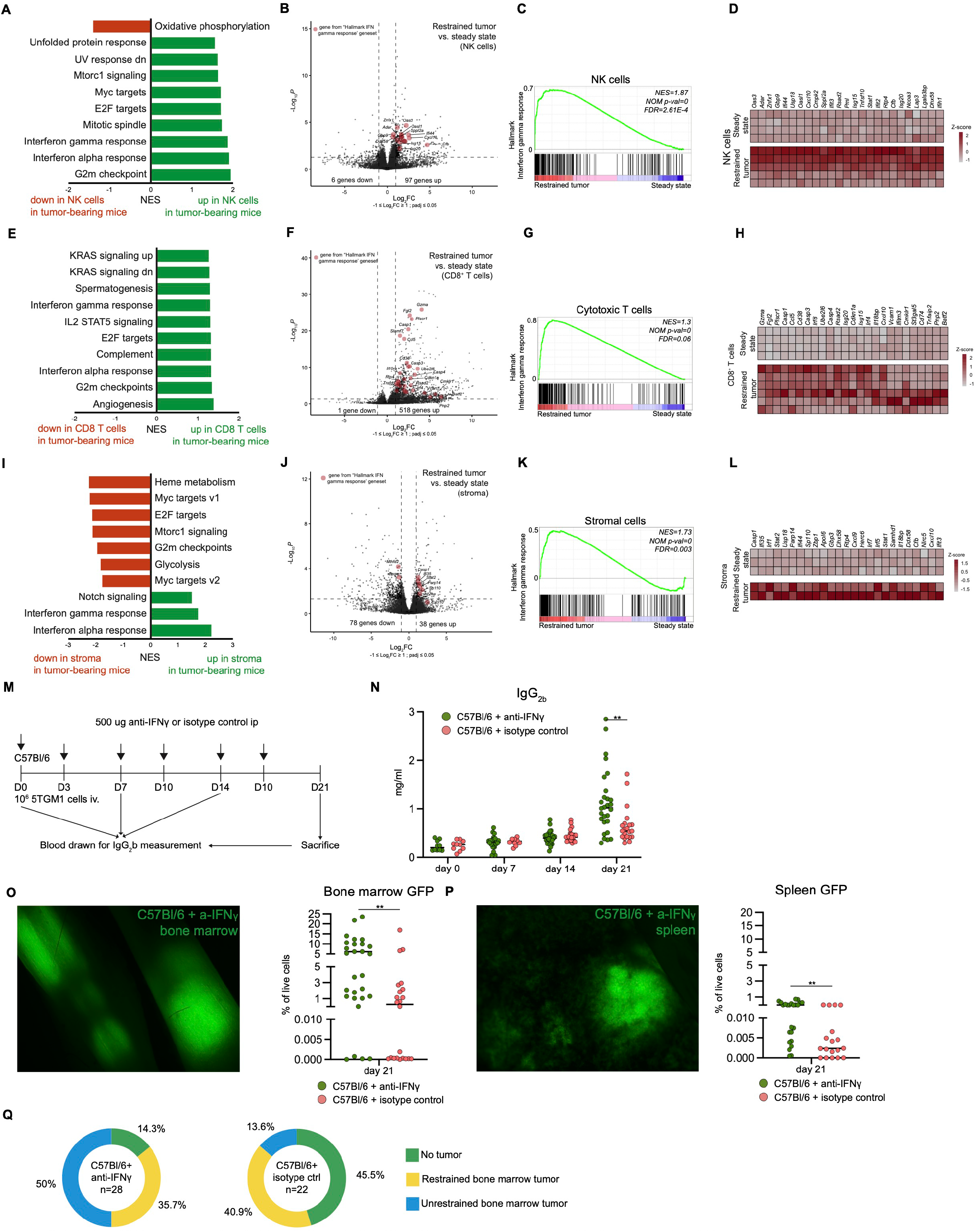
Neutralization of IFNy increases myeloma incidence and severity. **A**, GSEA of RNA sequencing from bone marrow NK cells showing top altered Hallmark pathways in bone marrow from mice with restrained tumors (n=5) compared to steady state (n=4). **B**, Volcano plot (*Log2FC cutoff±1; Padj* < *0.05 was considered significant)*, **C**, GSEAenrichment plot and **D**, Heat map showing selected top differential genes from the Hallmark ‘Interferon gamma response’ gene set in bone marrow NK cells. E, GSEA of RNA sequencing from bone marrow CDB T cells showing top altered Hallmark pathways from mice with restrained tumors (n=6) compared to steady state (n=4). F, Volcano plot *{Log2FC cutoff ±1; Padj* < *0.05 was considered significant)*, **G**, GSEA enrichment plot and **H**, heat map showing selected top differential genes from the Hallmark ‘Interferon gamma response’ gene set in bone marrow cytotoxic T cells. I, GSEA of RNA sequencing from bone marrow stromal cells showing top altered Hallmark pathways from mice with restrained tumor (n=2) compared to steady state (n=3). **J**, Volcano plot *(Log2FC cutoff ±1; Padj* < *0.05 was considered significant)*, **K**, GSEA enrichment plot and L, heat map showing selected top differential genes from the Hallmark ‘Interferon gammaresponse’ gene set in bone marrow stromal cells. **M**, Experimental setup of IFNy neutralization experiment. D: day; iv: intravenous injection. N, lgG2b levels in plasma after 5TGM1 transfer. 0_1_ Representative image of femurs from mice receiving anti-IfNy (left, 5x magnification) and percentage of bone marrow GFP+ tumors (right). **P**, Representative image of spleen from anti-lFNy-treated mice (left, 5x magnification) and percentage of splenic GFP• cells (right). **Q**, Donut graphs depicting myeloma growth patterns in C57BI/6 treated with anti-IfNy (left, n=28) and C57BI/6 treated with isotype control (right, n=22) mice. Green, no tumor; yellow, restrained bone marrow tumor; blue, unrestrained bone marrow tumor. Data are from at least 3 independent experiments. Line indicates median of values. Statistical significance in A-L was assessed using the Wald test followed by a Benjamini-Hochberg correction; and in N-P using Mann-Whitney U tests (ns, *p>0.05;* * *p<0.05;* ** *p*<0.01; \*\*\**p*<0.001; \*\*\*\**p*<0.0001).

Similar to T and NK cells, the ‘Interferon gamma response’ and ‘Interferon alpha response’ pathways were significantly enriched in C57Bl/6 day 21 stromal cells compared to steady state (**Fig5I-L, Supplementary Fig6E-G**).

As transcriptomic data from all these BM cell types indicated active IFNγ signaling in mice with myeloma, we next set out to test the contribution of this pathway to anti-myeloma immune responses following 5TGM1 transfer. Mice received IFNγ neutralizing antibodies or isotype control intraperitoneally on day 0 (prior to 5TGM1 transfer) and on days 3, 7, 10, 14, and 17, before being sacrificed on day 21 (**Fig5M**). Following IFNγ-neutralization, serum IgG_2b_ levels at day 21 were significantly increased compared to isotype controls (**Fig5N**). This correlated with increased BM tumor burden (**Fig5O**) and splenic tumor load (**Fig5P**). Importantly, tumor incidence was also increased after IFNγ neutralization (85.7%, 24 out of 28 vs 54.5%, 12 out of 22, p=0.0253) (**Fig5Q, Supplementary Table III**). This latter finding suggests that absence of tumor development in part of the C57BL/6 mice in our previous experiments was the result of immune-mediated eradication of the transferred cells. The increases in percentages of unrestrained BM tumor (58.3%, 14 out of 24 vs 25%, 3 out of 12) and splenic involvement (66.7%, 16 out of 24 vs 33.3%, 4 out of 12) did not reach statistical significance (p=0.0815 and p=0.083, respectively).

Together these results identify IFNγ as a relevant mediator of successful anti-myeloma immune responses in the 5TGM1 transfer model.

## Discussion

The tumor immune microenvironment is increasingly recognized as an important determinant of tumor progression and therapy response. In this work we analyzed the immune response to murine 5TGM1 MM cells in a well-known transfer model of the disease. To our knowledge, we provide the first comprehensive comparison of anti-5TGM1 immune responses in BM immune environments of C57Bl/6 versus KaLwRij mice during myeloma development.

While myeloma develops in 100% of injected KaLwRij mice, the parental C57Bl/6 strain is less permissive to tumor development (15). In contrast, RAG-2^-/-^ mice (lacking T and B cells) and NSG mice (lacking T, B, and NK cells) develop myeloma after inoculation with 5TGM1 cells (15,34), highlighting the importance of immune cells for preventing tumor outgrowth after transfer of syngeneic 5TGM1 cells. Interestingly, myeloma did not develop after transfer of 5TGM1 cells in nude athymic mice lacking mature T cells (13,15). Stromal cells may also influence tumor take as co-inoculation of 5TGM1 cells with BM stromal cells isolated from tumor-bearing KaLwRij mice was sufficient to allow tumor outgrowth in C57Bl/6 mice (35). Whether this was associated with immune modulation has not been determined.

We hypothesized that the difference in disease onset and severity between the two strains could be due to differential immune control of the transferred cells. Already at steady state, KaLwRij mice had a lower number of mature BM NK cells, raising the possibility of an impaired first response to tumor cells in these mice. In MM patients, malignant plasma cells have been shown to secrete regulatory cytokines that impair NK function (36), and NK cell exhaustion has been associated with disease progression (37). In the 5TGM1 system, activation of cytotoxic NK cells after tumor transfer occurs in both C57Bl/6 and KaLwRij mice, yet upon unrestrained tumor growth numbers of activated NK cells decline. This could be the result of either impaired expansion, or reduced recruitment into the BM from the circulation. Indeed, in line with our results, tumor-bearing KaLwRij mice develop reduced chemokine-mediated recruitment of NK cells (38). Altered recruitment could also explain the differences seen in the absolute number of CD8^+^ cytotoxic T cells of mice with unrestrained tumor growth. However, CD8 T cells also had an impaired activation status, maintaining expression of L-Selectin^+^ and absence of PD1. Induction of PD-1 expression is associated with exhaustion (39) but also identifies tumor-specific cytotoxic T cells in solid tumors (40). Our observation of minimal numbers of PD-1-expressing CD8 T cells in KaLwRij seems to be divergent from recent data by Fernandez-Poma et al. (41) where PD-1^+^ cyototoxic T cells were identified and sorted from tumor-bearing KaLwRij BM. This discrepancy might be due to experimental differences, as 5×10^6^ 5TGM1-GFP cells were injected, mice were analyzed 5-7 weeks after tumor transfer, and at this timepoint BM GFP% and IgG_2_b levels were much lower compared to our findings 3 weeks after tumor transfer (41). A more detailed analysis of CD8 T cells under various tumor conditions will be required to reconcile these findings.

In scRNA-seq analyses of human myeloma BM samples IFN-responsive cell subsets have been consistently identified (6-8). IFNγ also emerged as a central regulator from our transcriptomic studies, and neutralization of IFNγ led to a significant increase in tumor burden, suggesting a central role for this cytokine in the anti-myeloma response. A similar role for IFNγ was suggested by transfer of Vk-Myc MM cells into IFNγ^-/-^ mice (42). Importantly, the increased tumor incidence after IFNγ neutralization suggests that this cytokine is involved in immunological tumor clearance and prevention of measurable tumor outgrowth in approximately 30% of C57Bl/6 mice. IFNγ can be secreted by many different cell types within the tumor microenvironment, including NK cells (43) and cytotoxic T cells (44), but also T_H_1 cells, T_H_17 cells, *γδ* T cells [reviewed in (45)]. Here we did not identify the source(s) of IFNγ in the BM. The potential effects of IFNγ in context of tumor immunity are broad, as all nucleated cells can express the receptor for IFNγ. Downstream effects include direct anti-tumor responses, activation of cytotoxic T cells (46) and NK cells (47), and recruitment of effector cytotoxic cells by induction of CXCL9, CXCL10 and CXCL11 in the tumor microenvironment (48).

In summary, we show that C57Bl/6 mice develop myeloma after transfer of 5TGM1 cells, following an initial period of immune-mediated and IFNγ-dependent tumor containment. The C57Bl/6 background makes this a valuable model for studying the BM immune microenvironment in response to myeloma cells. Importantly, our data indicate that even though KaLwRij mice harbor a full immune system, immune responses in this strain differ from those in the parental C57Bl/6 strain. This will be an important consideration in studies evaluating anti-myeloma immune responses in these experimental models. Finally, as most immunologically relevant transgenic and knockout mice are in the C57Bl/6 genetic background, the ability to use C57Bl/6 mice for myeloma studies will expand the possibilities of manipulating the BM immune microenvironment in order to better understand its role in progression and therapy response of MM.

## Author contribution

T.C. and Z.K. conceptualized the study. T.C. and Z.K. were responsible for study methodology. Z.K., M.D.J., N.P., S.T., A.C.F. and C.D.H. were responsible for study investigation. Z.K., M.D.J., N.P., S.T., A.C.F., R.H., M.S., A.B., P.S. and T.C. analyzed the data. L.B. provided resources for the study. Z.K., M.D.J., R.H. and M.S. curated the data. Z.K. and M.D.J. contributed to data visualization. Z.K. and T.C. wrote the original draft, and all authors reviewed and edited the final manuscript. Z.K. acquired funding. T.C. supervised the study.

## Notes

**Financial Support** This project received funding from the European Union’s Horizon 2020 research and innovation program under the Marie Skłodowska-Curie grant agreement no. 707404 (to Z.K.) The European Commission is not responsible for any use that may be made of the information it contains.

**Competing interests** A.B. consults for BMS/Celgene, Janssen, Amgen and Sanofi. P.S. is on the advisory board for Amgen, BMS/Celgene, Janssen, Seagen and Pfizer and receives research support from Janssen, BMS/Celgene and Karyopharm.

### Competing Interest Statement

Competing interests
A.B. consults for BMS/Celgene, Janssen, Amgen and Sanofi. P.S. is on the advisory board for Amgen, BMS/Celgene, Janssen, Seagen and Pfizer and receives research support from Janssen, BMS/Celgene and Karyopharm.

