## Supplementary information for "Immune-mediated tumor control in the 5TGM1 transfer model of multiple myeloma"

**Supplementary Figure 1.**

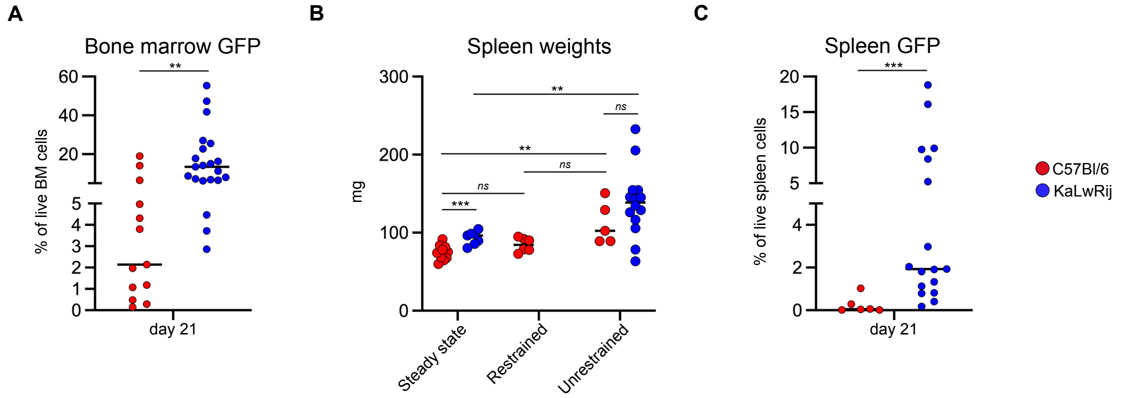


**Supplementary Figure 1 (related to Figure 1). A**, Percentage of GFP⁺ tumor cells in bone marrows of tumor-bearing mice. **B**, Spleen weights. **C**, Percentage of splenic GFP⁺ tumor cells in mice with splenic dissemination. Data are from at least 3 independent experiments. Line indicates median of values. Statistical significance was assessed using Mann-Whitney *U* tests (ns, *p*>0.05; *, *p*<0.05; **, *p*<0.01; ***, *p*<0.001;

**Supplementary Figure 2.**

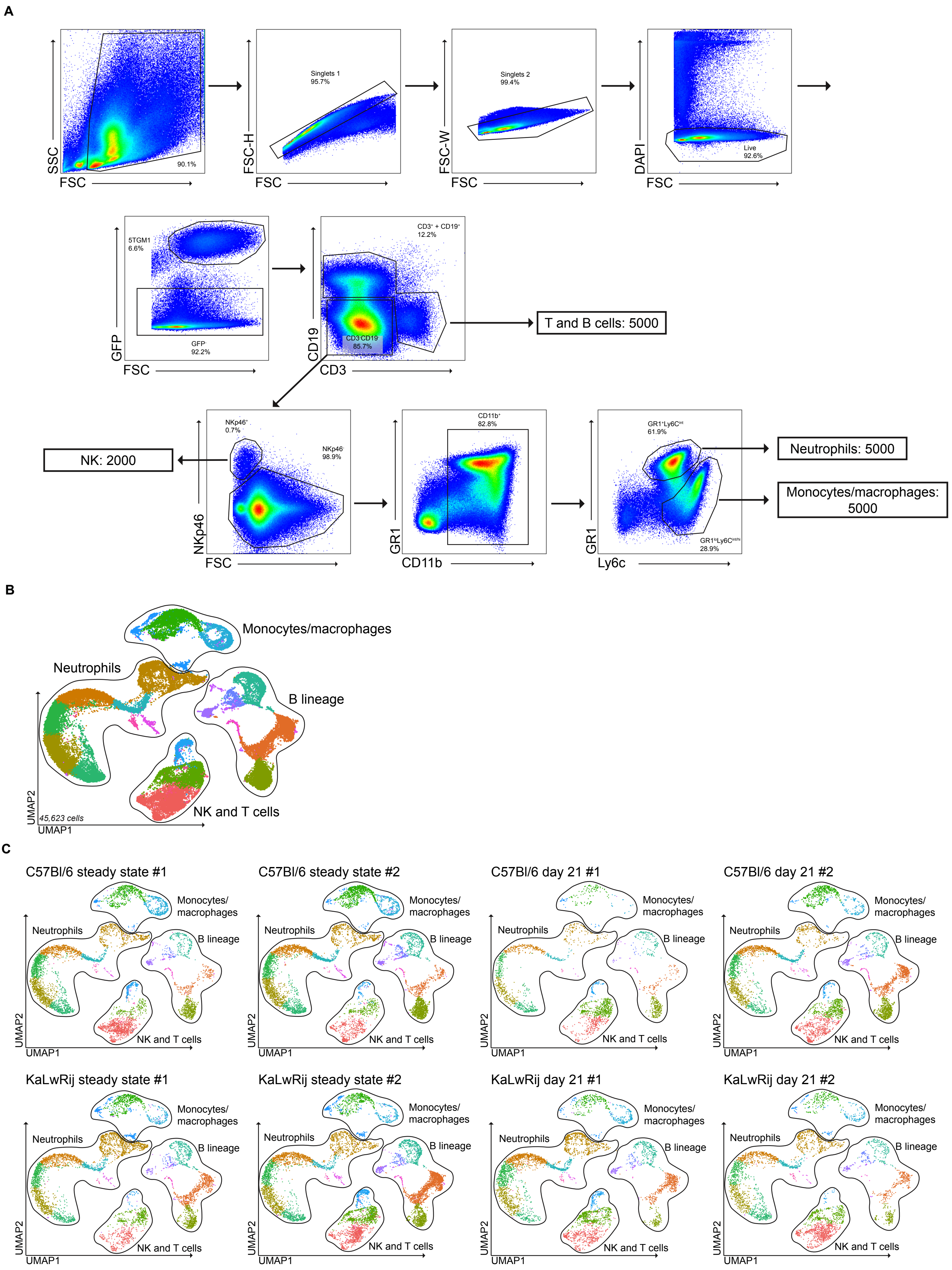


**Supplementary Figure 2 (related to Figure 2). A,** Sort strategy for single-cell sequencing of bone marrow NK, T, B, monocyte/macrophage and neutrophil populations. **B,** UMAP of all 45,623 cells recovered from 8 mice. Colors indicate the 22 clusters obtained at resolution = 0.4. **C,** UMAPs of individual mice, subsetted from the full object. Colors indicate the the 22 clusters obtained at resolution = 0.4.

**Supplementary Figure 3.**

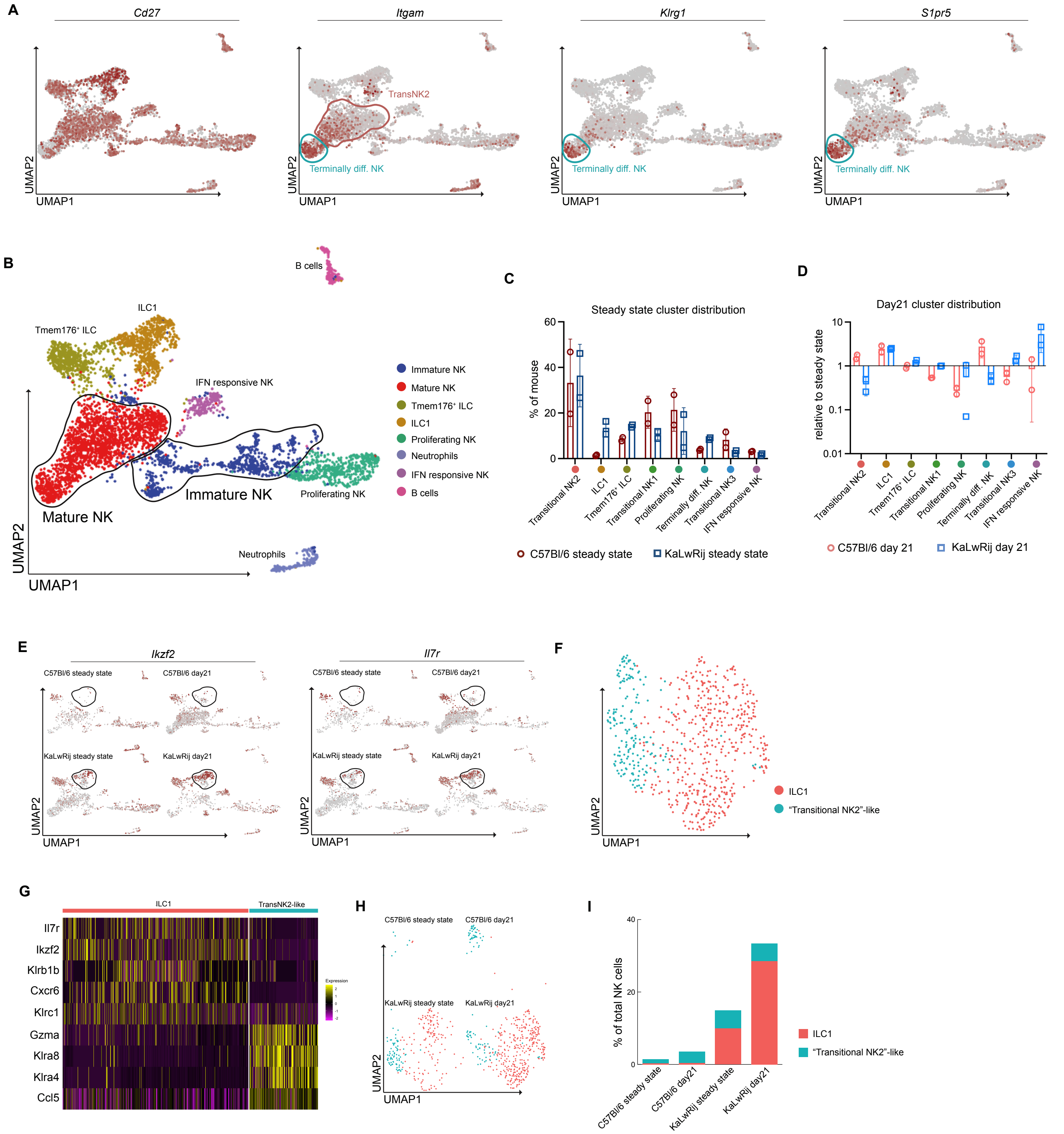


**Supplementary Figure 3 (related to Figure 2). A,** Feature plots of genes associated with NK cell maturation stages; circled areas refer to indicated clusters colored using respective UMAP colors. **B,** UMAP with appropriate clusters grouped into 'Immature' and 'Mature' NK cells. **C,** Bar graphs showing frequencies of individual clusters under steady state conditions. **D,** Bar graphs showing frequencies of individual clusters in tumor-bearing mice compared to steady state. **E,** Feature plots split per condition showing Ikzf2 and Il7r transcription; circled area indicates 'ILC1' cluster. **F,** UMAP of subsetted and reanalyzed 'ILC1' cluster. **G,** Heat map showing top selected differentially expressed genes of subsetted and reanalyzed 'ILC1' cluster. **H,** Split UMAPs of subsetted and reanalyzed 'ILC1' cluster. **I,** Column charts depicting relative number of cells within 'ILC1' and 'Transitional NK2-like' clusters as % of all NK cells.

**Supplementary Figure 4.**

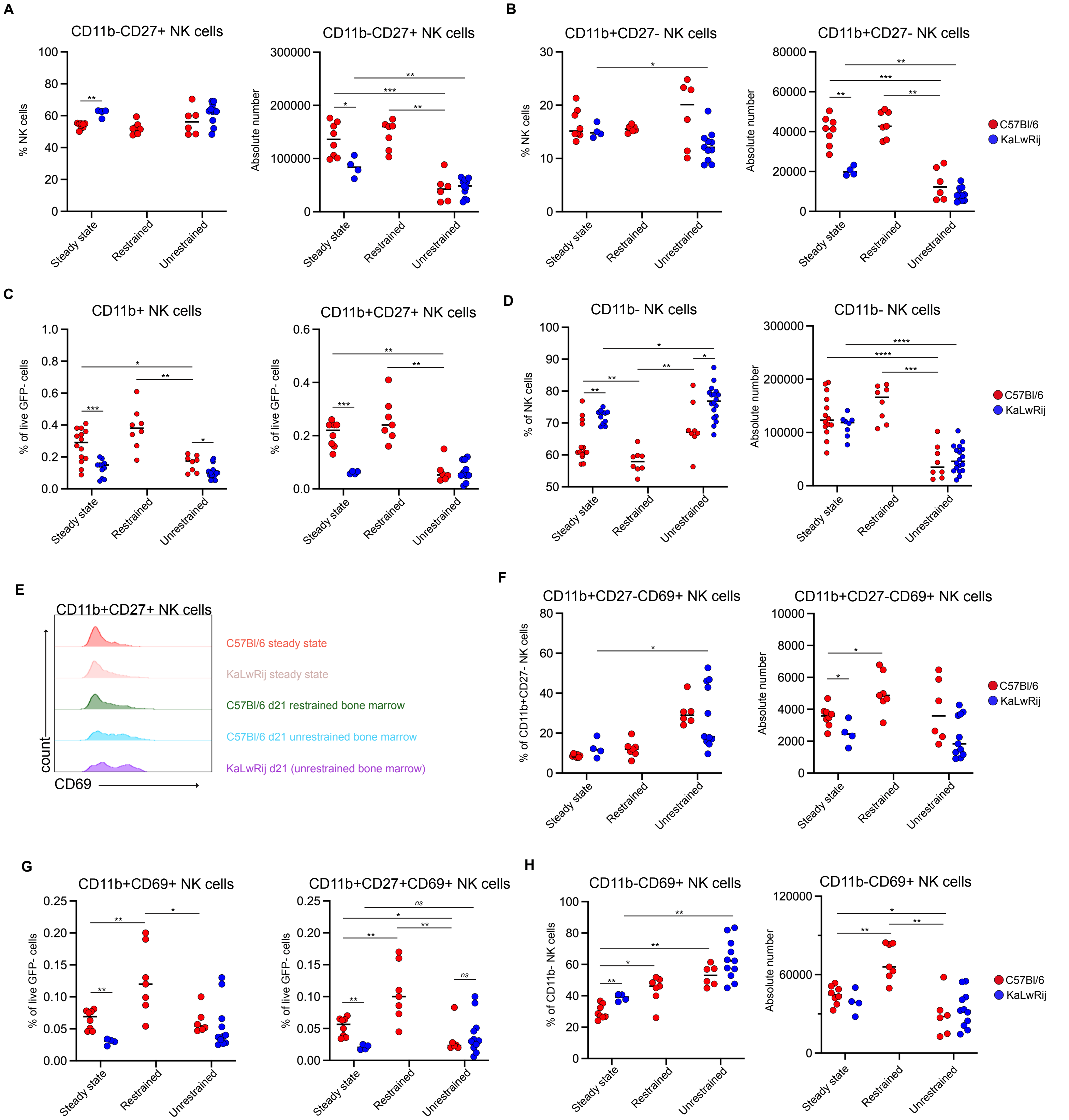


**Supplementary Figure 4 (related to Figure 3). A,** Relative (left) and absolute (right) number of bone marrow CD11b$^-$CD27$^+$ NK cells. **B,** Relative (left) and absolute (right) number of bone marrow CD11b$^+$CD27$^-$ NK cells. **C,** Relative numbers of CD11b$^+$ NK cells (left) and CD11b$^+$CD27$^+$ NK cells (right), shown as percentage of GFP$^-$ live cells. **D,** Relative (left) and absolute (right) number of CD11b$^-$ bone marrow NK cells. **E,** Histogram from representative samples showing CD69 expression on CD11b$^+$CD27$^+$ NK cells. **F,** Relative (left) and absolute (right) number of bone marrow CD11b$^+$CD27$^-$CD69$^+$ NK cells. **G,** Relative numbers of bone marrow CD11b$^+$CD69$^+$ NK cells (left) and CD11b$^+$CD27$^+$CD69$^+$ NK cells (right), shown as percentage of GFP$^-$ live cells. **H,** Relative (left) and absolute (right) number of CD11b$^-$CD69$^+$ bone marrow NK cells. Data are from at least 3 independent experiments. Line indicates median of values. Statistical significance was assessed using Mann-Whitney *U* tests (ns, $p>0.05$; *, $p<0.05$; **, $p<0.01$; ***, $p<0.001$; ****, $p<0.0001$).

**Supplementary Figure 5.**

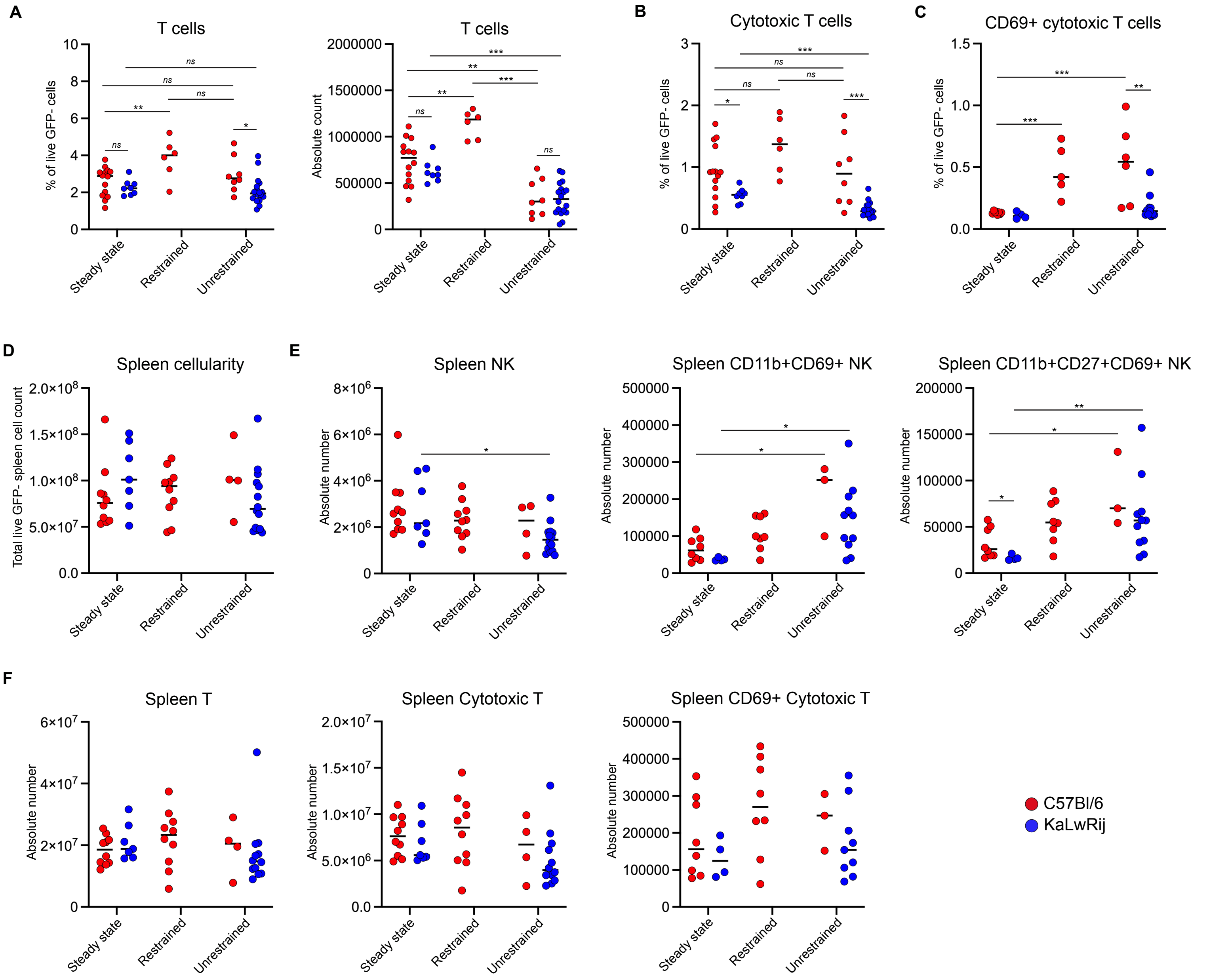


**Supplementary Figure 5 (related to Figure 4). A,** Relative (left) and absolute (right) number of bone marrow T cells. **B,** Bone marrow cytotoxic T cells shown as percentage of live $GFP^-$ bone marrow cells. **C,** Bone marrow $CD69^+$ cytotoxic T cells shown as percentage of $GFP^-$ live cells. **D,** Absolute number of total GFP- live spleen cells. **E,** Absolute number of spleen NK cells (left), spleen $CD11b^+CD69^+$ NK cells (middle) and spleen $CD11b^+CD27^+CD69^+$ NK cells (right). **F,** Absolute number of spleen T cells (left), spleen cytotoxic T (middle), and spleen $CD69^+$ cytotoxic T cells (right). Data are from at least 3 independent experiments. Line indicates median of values. Statistical significance was assessed using Mann-Whitney *U* tests (ns, $p>0.05$; *, $p<0.05$; **, $p<0.01$; ***, $p<0.001$; ****, $p<0.0001$).

**Supplementary Figure 6.**

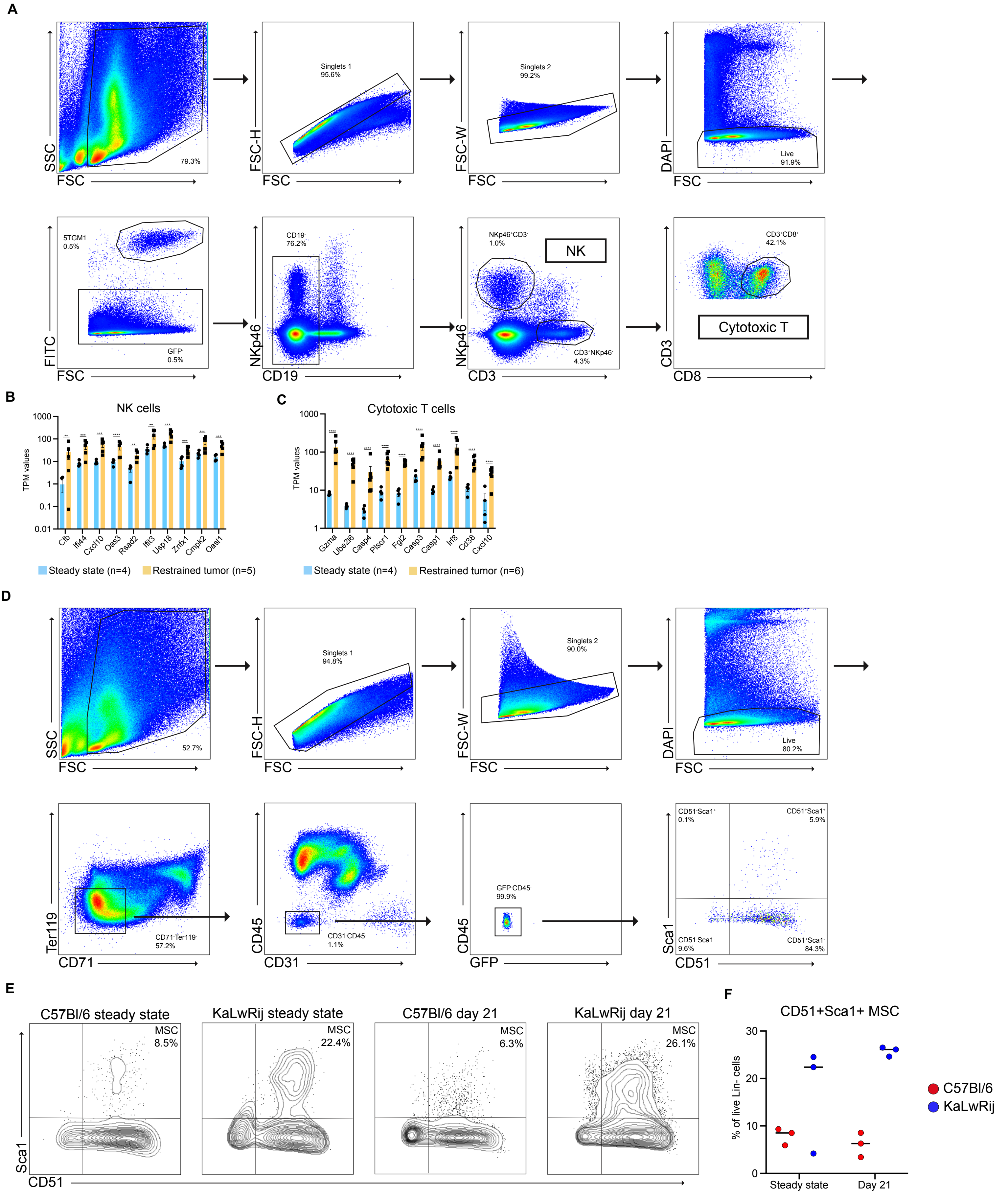


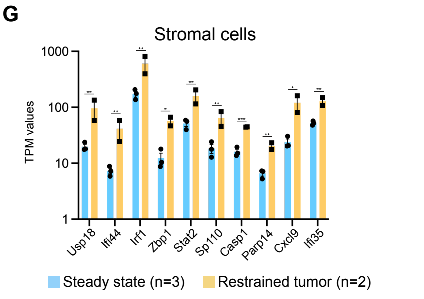


**Supplementary Figure 6 (related to Figure 5). A,** Sort strategy for bone marrow NK and cytotoxic T cells. **B,** Bar graph showing selected top differential genes from the Hallmark 'Interferon gamma response' gene set in bone marrow NK cells. **C,** Bar graph showing selected top differential genes from the Hallmark 'Interferon gamma response' gene set in bone marrow cytotoxic T cells. **D,** Sort strategy for bone marrow stroma cells. **E,** Representative flow cytometric plots of bone marrow stromal cells. **F,** Percentage of CD51$^{+}$Sca1$^{+}$ MSCs. **G,** Bar graph showing selected top differential genes from the Hallmark 'Interferon gamma response' gene set in bone marrow stromal cells. Data in B, C, and G are shown as mean ± SEM. Statistical significance in B, C, and G was assessed using the Wald test followed by a Benjamini-Hochberg correction (ns, $p>0.05$; *, $p<0.05$; **, $p<0.01$; ***, $p<0.001$; ****, $p<0.0001$).

| Supplementary Table I. | Total | Tumor | | Restrained BM tumor | Unrestrained BM tumor | | Splenic involvement | |
|---|---|---|---|---|---|---|---|---|
| KaLwRij | 21 | 100% (21/21) | ***p=0.0014* | 14.3% (3/21) | 85.7% (18/21) | ****p=0.0006* | 100% (21/21) | ****p=0.0003* |
| C57Bl/6 at day 21 | 22 | 59.1% (13/22) | *ns, p>0.9999* | 76.9% (10/13) | 23.1% (3/13) | *ns, p=0.0623* | 46.2% (6/13) | *ns, p=0.3742* |
| C57Bl/6 long term | 12 | 58.3% (7/12) | | 28.6% (2/7) | 71.4% (5/7) | | 71.4% (5/7) | |

**Supplementary Table I.** Tumor incidence, bone marrow growth pattern, and splenic involvement in KaLwRij mice at day 21 and C57Bl/6 mice at day 21 or after (long-term experiment). Significance was assessed using Fisher's exact test.

| Supplementary Table II. | BM GFP % | Plasma IgG2b mg/ml | Total recovered cell count | NK cells | T cells |
|---|---|---|---|---|---|
| C57Bl/6 steady state #1 | - | - | 6389 | 336 | 924 |
| C57Bl/6 steady state #2 | - | - | 7600 | 312 | 470 |
| C57Bl/6 day 21 #1 | 0.32% | 2.07 mg/ml | 2451 | 794 | 112 |
| C57Bl/6 day 21 #2 | 1.97% | 0.61 mg/ml | 5357 | 667 | 426 |
| KaLwRij steady state #1 | - | - | 6145 | 373 | 289 |
| KaLwRij steady state #2 | - | - | 8373 | 1097 | 594 |
| KaLwRij day 21 #1 | 55.3% | 7.53 mg/ml | 5144 | 723 | 405 |
| KaLwRij day 21 #2 | 16.3% | 7.18 mg/ml | 4164 | 472 | 603 |

**Supplementary Table II.** Characteristics of mice used for single-cell sequencing (bone marrow GFP%, plasma $IgG_{2b}$ levels, number of recovered cells, and number of NK cells and T cells).

| Supplementary Table III. | Total | Tumor | | Restrained BM tumor | Unrestrained BM tumor | | Splenic involvement | |
|---|---|---|---|---|---|---|---|---|
| C57Bl/6 + anti-IFN$\gamma$ | 28 | 85.7% (24/28) | **p=0.0253* | 41.7% (10/24) | 58.3% (14/24) | *ns, p=0.0815* | 66.7% (16/24) | *ns, p=0.083* |
| C57Bl/6 +isotype control | 22 | 54.5% (12/22) | | 75% (9/12) | 25% (3/12) | | 33.3% (4/12) | |

**Supplementary Table III.** Tumor incidence, bone marrow growth pattern, and splenic involvement in C57Bl/6 mice treated with anti-IFNg or isotype control. Significance was assessed using Fisher's exact test.
